# Insight into the evolution of symbiosis in the *Cupriavidus* genus: high conserved symbiotic island and a patchy phylogenetic distribution

**DOI:** 10.64898/2026.03.04.709584

**Authors:** Melisa Magallanes Alba, Cecilia Rodríguez-Esperón, José Sotelo-Silveira, Elena Fabiano, Andrés Iriarte, Raúl Platero

## Abstract

Currently, there are three recognized rhizobial genera belonging to the beta branch of the proteobacteria; *Trinickia, Paraburkholderia,* and *Cupriavidus.* These beta-rhizobia have been found associated with legume species mainly within the *Mimosoideae* and *Papillonoideae.* Most diversity, evolutionary, and functional studies have focused on Paraburkholderia, whereas few have addressed the diversity and evolution of symbiosis in the *Cupriavidus* genus. The present work aimed to provide an actual view of the symbiotic *Cupriavidus* diversity and to analyse the origin and evolution of their symbiotic genes.

Using whole-genome information for phylogenetic reconstruction, we showed that the described symbiotic Cupriavidus strains belong to five distinct lineages, although they are intermixed with non-symbiotic species. The high synteny and sequence conservation of symbiotic genes suggest a common origin of acquisition for all rhizobial *Cupriavidus* described so far. However, we observed very low sequence conservation among (mega)plasmids carrying the symbiotic island, excluding the existence of a conserved symbiotic plasmid within beta-rhizobia.

We can conclude that up to now there are five rhizobial species within the *Cupriavidus* genus, and we predict the description of new symbiotic species in the near future.

## Introduction

Rhizobia are physiological and genetically diverse soil bacteria able to symbiotically interact with legume plants (Granada Agudelo et al., 2023). During these interactions, novel organs known as nodules are formed. Within these nodules, bacteria are housed intracellularly and express the genes necessary for biological nitrogen fixation (BNF). This mutualistic symbiosis involves the plant nourishing the nodule bacteria, which in turn provide the plant with sufficient nitrogen for its complete development (Chiurazzi et al., 2025). Currently, there are about 14 recognized genera of rhizobia distributed across the alpha and beta subgroups of the family Proteobacteria (Andrews & Andrews, 2017).

First described in 2001 (W. M. Chen et al., 2001; Moulin et al., 2001), rhizobia belonging to the *Burkholderiaceae*, the so-called Beta-rhizobia, have been found associated with numerous legume plants (Bellés-Sancho et al., 2023). Within *Burkholderiaceae,* three genera of beta-rhizobia have been described; *Trinickia, Paraburkholderia,* and *Cupriavidus* (Coenye, 2014; Estrada-de Los Santos et al., 2018). The *Burkholderia* genus is composed of a growing number of species that can be classified into two main groups according to their legume preference; one group was found associated with legume members of the *Papillonoideae* in the Fynbos region in South Africa, while the second forms a bigger group and is associated with legumes in the tribe *Mimoseae*. Accordingly, each group has a particular symbiotic gene set and is not able to form symbiotic associations with legume partners of the other group (Lemaire et al., 2016).

In contrast, beta-rhizobia *Cupriavidus* are believed to be much less diverse than *Burkholderia*, since they have only been found forming effective symbiotic associations with legume members within the *Mimoseae* subfamily (Bellés-Sancho et al., 2023; Platero et al., 2016).

*Cupriavidus* is a large genus within the β subclass of proteobacteria, best known for its ability to degrade xenobiotic compounds (Cuadrado et al., 2010; Lykidis et al., 2010; Pérez-Pantoja et al., 2008), its high metal resistance (Janssen et al., 2010), and as a model organism for poly-β-hydroxybutyrate metabolism (PHB) (Nygaard et al., 2021). Recent studies have expanded the ecological and metabolic scope of the genus, showing that *Cupriavidus ulmosensis* sp. can also colonize extreme environments and perform carbon monoxide oxidation (Dawson et al., 2025). However, the description in 2001 of a novel *Cupriavidus sp.* isolated in Taiwan from nodules of an exotic legume plant, *Mimosa pudica,* opened new horizons for this genus (W. M. Chen et al., 2001). From this initial report, *C. taiwanensis* strains were isolated from many exotic invasive *Mimosa* species in different parts of the world, but were also associated with Mimosa *spp.* within its native range in the Americas (Andrus et al., 2012). Barrett and Parker first reported the presence of *Cupriavidus* as a symbiont of *Mimosa* sp. growing in its native habitat for *M. pudica* and *M. pigra* populations from Costa Rica (Barrett & Parker, 2006). A year later, in Texas, USA, a population of *Mimosa asperata* (a close relative of *M. pigra*) whose nodules were dominated by *Cupriavidus sp.* symbionts was found. Interestingly, analysis of the *nodA* and *nifH* sequences of these isolates showed high homology with the symbiotic genes of the previously described Taiwan isolates, suggesting that symbiotic genes for both groups could derive from a recent common ancestor (Andam et al., 2007a).

In contrast to these findings, many surveys have been performed focusing on rhizobial diversity of *Mimosa* and related genera in different Brazilian biomes, including the Cerrado (savanna), Caatinga (xeric shrubland), and Pantanal (wetland) (Bontemps et al., 2010; Dos Reis Jr et al., 2010), Amazônia and Mata Atlântica (W.-M. Chen et al., 2005). In all these studies, the majority of the symbionts belonged to the *Paraburkholderia* genus. However, Da Silva and collaborators were able to isolate *Cupriavidus* strains from nodules of *Phaseolus vulgaris* grown in soil from Minas Gerais, Brazil. Surprisingly, phylogenetic and polyphasic analyses indicated that these strains belong to the *Cupriavidus necator* species rather than to the described beta-rhizobia *C. taiwanensis* (Da Silva et al., 2012).

Adding to this geographical and host expansion, recent studies in Mexico have revealed even greater symbiotic diversity within the genus. Several novel species have been proposed, such as *C. phytorum* (associated with *Mimosa diplotricha* root nodules) and *C. consociatus* (isolated from *Arachis* sp. and *Leucaena* sp. nodules). Moreover, Tapia-García and collaborators (Tapia-García et al., 2026) recently described the new species *Cupriavidus phytohabitans*, isolated from *Phaseolus vulgaris* trap plants inoculated with rhizospheric soil from wild *Acacia* sp. in Veracruz. Notably, genomic comparisons demonstrated that the AMP6 strain, previously isolated from Mimosa asperata in Texas, also belongs to this newly defined *C. phytohabitans* species.

Also, *Cupriavidus sp.* strains were isolated and characterized from nodules of wild populations of the Mimosoideae tree *Parapiptadenia rigida* in Uruguay (Taulé et al., 2012). Phylogenetic analyses showed that these beta-rhizobia strains are also close relatives to *C. necator*, probably belonging to the same group as the Brazilian isolated strains. Interestingly, these *Cupriavidus sp.* were shown to be able to nodulate and promote plant growth both in their original host and in various members of the *Mimosa* genus, including *M. pudica* (Taulé et al., 2012). These findings motivated us to conduct a country-wide survey of *Mimosa* symbionts, finding that a high proportion of the sampled species harbours beta rhizobia belonging to the *Cupriavidus* genus. Along with these *C. necator* isolates, we identified a second cluster of *Cupriavidus* strains that did not group with any known symbiotic *Cupriavidus* species and could represent a new beta-rhizobia species (Platero et al., 2016).

Genome sequencing of the *Cupriavidus taiwanensis* type strain LMG19424 revealed the presence of a 35 kb symbiotic island, where the *nod*, *nif*, and *fix* genes are organized into a few operons. This compact arrangement of the abundance, together with insertion sequences and transposase genes, has been interpreted as evidence of a recent evolutionary acquisition of symbiosis-related traits in this lineage (Amadou et al., 2008). Symbiotic gene analysis revealed that, like in most alpha-rhizobia, *nod* and *nif* genes were present, suggesting a common strategy adopted by both alpha and beta-rhizobia to establish successful symbiotic interactions with legumes (Black et al., 2012).

Surprisingly, comparative genomic analyses performed with available rhizobial genomes at that time revealed that less than 10% of the genes were shared by all rhizobia (Amadou et al., 2008; Garrido-Oter et al., 2018). This study clearly showed that no set of genes is both common and exclusive to all rhizobia, highlighting the absence of a unique genetic program universally responsible for symbiosis with legumes (Amadou et al., 2008; Tian et al., 2012).

In addition, our group sequenced and annotated the genome of *Cupriavidus* sp. strain UYMMa02A, isolated from *Mimosa magentea* in Uruguay. Previous studies on this strain demonstrated its capability to nodulate Mimosa species efficiently even in the absence of typical flavonoid induction (Rodríguez-Esperón et al., 2023). This strain does not cluster with any previously known beta-rhizobial *Cupriavidus*, representing a novel lineage within this group (Iriarte et al., 2016).

In the present work, we have exploited the available genomic data from different *Cupriavidus* species to study the evolutionary history of symbiosis in the genus. We analyse the particularities of each symbiont, the organization of the symbiotic cluster, and that of the symbiotic plasmid. Our results support the existence of novel symbiotic lineages within South America, leading to the description of *C. ariasiae* sp. nov. Furthermore, comparative analysis of the symbiotic plasmid reveals a mosaic evolutionary pattern: while the symbiotic island remains highly conserved across the genus, the plasmid backbone exhibits significant variability. This contrast suggests that, beyond vertical inheritance in *C. taiwanensis*, horizontal gene transfer and plasmid remodeling play crucial roles in the dissemination of symbiosis in *Cupriavidus*.

## MATERIALS AND METHODS

### Genome sequences for comparative genomics

Available genomes of the *Cupriavidus* genus were downloaded from the NCBI database (Sayers et al., 2022) via NCBI Datasets command-line tools (O’Leary et al., 2024) (November, 2025). The obtained genomes were filtered based on their assembly level. However, to ensure that no symbiotic lineage was inadvertently excluded by this quality-based filtering, we performed an exhaustive BLASTn search against the entire initial dataset using the complete symbiotic island of *C. taiwanensis* LMG19424^T^ as a query. Any genome displaying significant homology coverage to the symbiotic island was retained and incorporated into the final dataset, regardless of its assembly rank. In addition, all available type strains were validated and incorporated into the dataset based on EzBioCloud taxonomic assignments (Chalita et al., 2024). The final datasets comprised 114 genomes, including three *Paraburkholderia* genomes incorporated as outgroups in the analysis (Table S1).

### Genome comparative analysis

To assess the taxonomic status of strain *Cupriavidus sp.*UYMMa02A and elucidate the genomic structure of the genus, a comprehensive All-vs-All comparative genomics analysis was conducted among *Cupriavidus* type strains and relevant isolates. Average Nucleotide Identity (ANI) was computed using two complementary algorithms: FastANI v1.3 (Jain et al., 2018) for rapid estimation, and ANIb (BLAST-based ANI) using PyANI v0.2.13 (Pritchard et al., 2016) for high-precision alignment. Subsequently, digital DNA–DNA hybridization (dDDH) values were estimated for the complete dataset using a Generalized Linear Model (GLM). This model transforms ANI values into predicted dDDH probabilities based on the empirical correlation established by (Meier-Kolthoff et al., 2013), allowing for rigorous species delimitation using the 70% dDDH threshold. Genomic distance matrices were visualized as hierarchical clustering heatmaps. Strains were clustered based on Euclidean distance and average linkage. To enhance interpretation, color scales were discretized to visually delimit taxonomic boundaries (ANI >96% and dDDH >70%), and species clusters were demarcated with boundaries.

Finally, a phylogenomic analysis was performed with PhyloPhlAn v3.1.1 (Asnicar et al., 2020), which utilizes 400 universal marker genes. The phylogenetic tree was inferred using Maximum Likelihood in IQ-TREE v2.4.0 (Minh et al., 2020), with the best-fit amino acid substitution model and branch support assessed via 1000 ultrafast bootstrap replicates. The resulting trees were visualized and annotated using iTOL (Letunic & Bork, 2024).

### Comparative genomics of nodulating *Cupriavidus* strains

To identify and compare symbiotic genes across all selected strains, we used as reference the symbiotic island sequence from *C. ariasiae* UYMMa02A to perform a protein-to-genome alignment using Miniprot v0.13 (Li, 2023) with default parameters and the standard bacterial genetic code (translation table 11). Each genome was independently aligned to the *C. ariasiae* symbiotic proteins, and resulting annotations were parsed to identify coding sequences corresponding to nodulation (*nod*), nitrogen fixation (*nif*, *fix*), and associated regulatory genes. We applied a conservative strategy by retaining only the top-scoring hit per genome for each gene. Since Miniprot performs splicing-aware alignment, it effectively handles frameshifts and internal stop codons while prioritizing full-length, high-confidence matches, ensuring that the best hit typically corresponds to the canonical ortholog (Li, 2023). Additional quality filters were applied to discard sequences with unexpected lengths or internal stop codons, and we confirmed the presence of conserved functional domains when applicable. For each gen, aligned loci corresponding to nodulation genes or conserved symbiotic markers were extracted in GFF3 format and used to retrieve protein-coding sequences with AGAT v1.4.0 (agat_sp_extract_sequences.pl), applying the --table 11 flag for correct codon translation (Jacques Dainat et al., 2024). Alignments for each orthologous group were generated with MAFFT v7.505 (Katoh & Standley, 2013) and concatenated using SEGUL v0.22.1 (Handika & Esselstyn, 2024). Phylogenetic trees were inferred using IQ-TREE v2.4.0 (Minh et al., 2020) under the best-fit substitution model (Kalyaanamoorthy et al., 2017), including support evaluation with 1,000 bootstrap replicates (-b 1000) and SH-aLRT tests (-alrt 1000) (Felsenstein, 1985). For those genomes in which the symbiotic island was identified, synteny analysis was subsequently performed using MCScanX (python) (Wang et al., 2012).

A BLASTn was performed to align the complete symbiotic plasmid of *Cupriavidus taiwanensis* LMG19424^T^ (query) against the Burkholderiaceae family (subject database). BLASTn hits were converted into BED format and merged with bedtools to compute non-overlapping coverage. For each subject we calculated: (i) total nucleotides aligned to the query plasmid (nt_total), (ii) nucleotides aligned within the coordinates of the symbiotic island (nt_in_island), (iii) nucleotides aligned outside the island (nt_out_island), and the derived percentages: pct_total (coverage of the plasmid), pct_isla_plasmid (coverage of the plasmid explained by the island), pct_out_plasmid (coverage of the plasmid outside the island), and pct_isla_relative (fraction of the island covered). These metrics are summarized in Table S3. The pipeline used to generate these metrics is available at https://github.com/Melisa-Magallanes/Plasmid-analysis.

## RESULTS AND DISCUSSION

### *Cupriavidus* genomes comparative analysis and phylogenetic diversity

An All-vs-All genomic comparison was performed to define the taxonomic position of the studied strains within the genus *Cupriavidus*. The resulting matrices (ANIb, FastANI, and estimated dDDH) revealed a clear genus-wide structure (Table S2), visualized in **Figure S1** as distinct species clusters delimited by high genomic identity values (ANI >96%; dDDH >70%). A focal subset comprising type strains and nodulating strains is shown in **Figure 1**. In the case of *C. ariasiae* UYMMA02A, the analysis confirmed its novelty. Its closest phylogenetic relative was *C. laharis*, but genomic similarity values were low (88.33% ANIb, 37.2% estimated dDDH), falling well below the species delineation thresholds (95–96% ANI, 70% dDDH). This discontinuity supports the description of *C. ariasiae* as a novel species. This taxonomic distinction is consistent with its unique physiological traits, distinct from *C. taiwanensis*, such as the divergent regulation of nodulation genes and specific proteomic responses recently described (Rodríguez-Esperón et al., 2023). Regarding *Cupriavidus* sp. UYPR2.512, the closest match was *C. necator* H850. However, comparison yielded values of 94.25% ANIb and 60.2% dDDH. When compared specifically to the type strain *C. necator* N-1^T^ (**Figure 1**), the strain yielded a similarly borderline ANIb value of 94.12%. These results place the strain in the speciation “grey zone”, consistently below the threshold for species identity, suggesting that *Cupriavidus* UYPR2.512 represents a distinct, diverging lineage within the *C. necator* complex.

**Figure 1.**
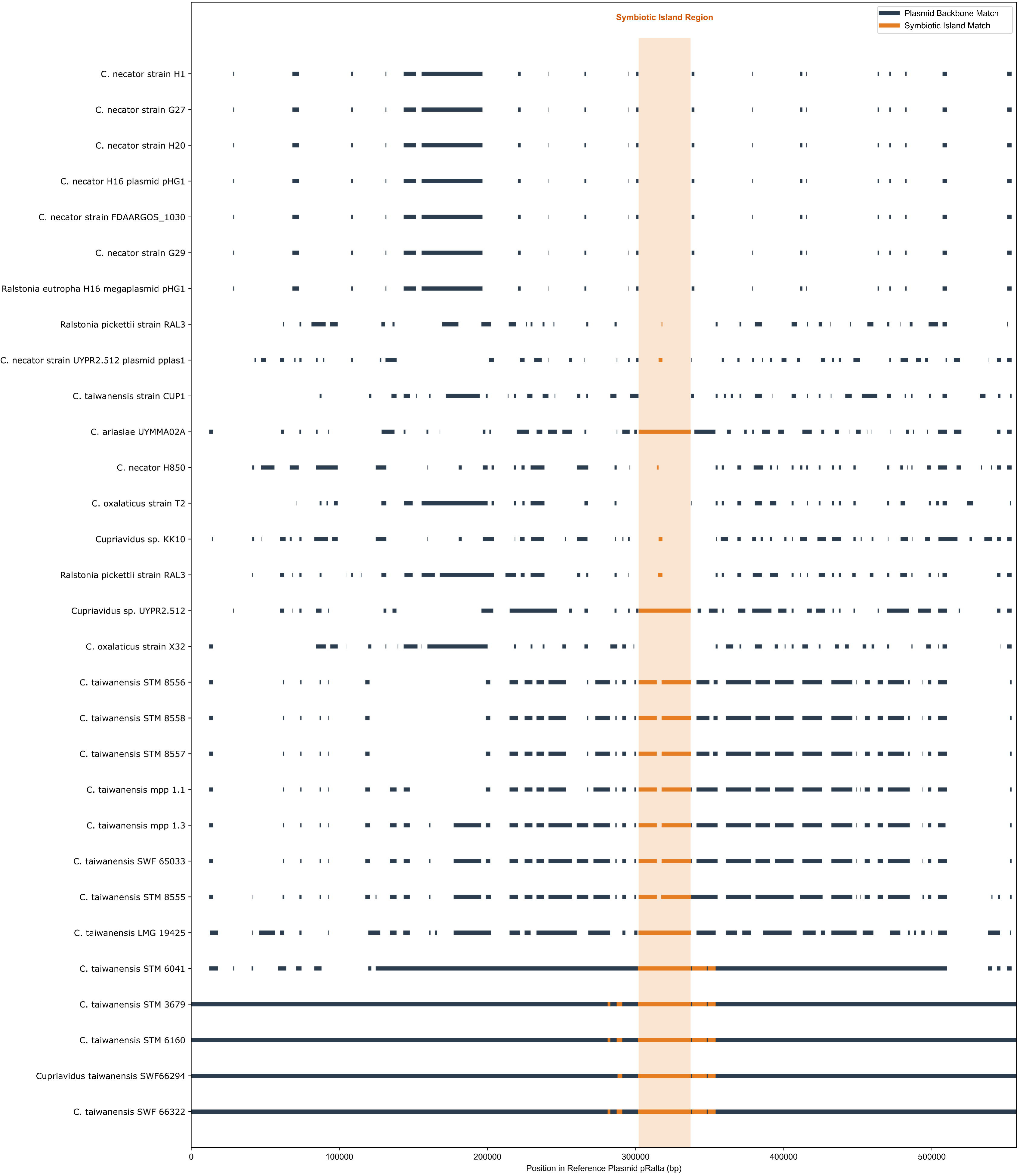
Genomic relatedness among *Cupriavidus* type strains and nodulating strains. The heatmap displays pairwise Average Nucleotide Identity (ANIb) values computed for a subset of genomes, including type strains and relevant nodulating strains, ordered according to the associated dendrogram, which groups strains based on genomic distance (Euclidean distance, average linkage). The color scale is discretized to reflect taxonomic boundaries: red indicates species-level identity (>96%), orange represents the speciation transition zone (95–96%), and blue/grey values (<90%) denote distinct species.

However, to clarify its taxonomic placement and evolutionary relationships, a phylogenomic analysis was performed using PhyloPhlAn 3.0, incorporating 111 *Cupriavidus* genomes and three *Paraburkholderia* species as outgroups, which revealed distinct evolutionary placements for the two reference strains. *Cupriavidus* sp. UYPR2.512 clustered within a Cupriavidus necator-*like* clade, including multiple type strains such as *C. necator* H850 and *C. necator* NBRC 102504. However, while *C. necator* H850 clustered tightly with the type strain *C. necator* N-1^T^, exhibiting a high ANIb value of 98.69% that unequivocally confirms its species status, the situation for *C. necator* NBRC 102504 was notably different. Although currently classified as *C*.

*necator*, strain NBRC 102504 shared only 94.81% ANIb with the type strain, falling below the accepted species demarcation threshold (95–96%). *Cupriavidus* sp. UYPR2.512 formed a phylogenetic clade with this misclassified strain, sharing similarly borderline ANIb values with the type strain (94.12%) and with *C. necator* H850 (94.25%). These findings suggest that both *Cupriavidus* sp. UYPR2.512 and *Cupriavidus necator* NBRC 102504 represent diverging lineages within the *C. necator* species complex, distinct from the *sensu stricto* cluster formed by strains *C. necator* N-1^T^ (Figure 2A and Table S2).

**Figure 2.**
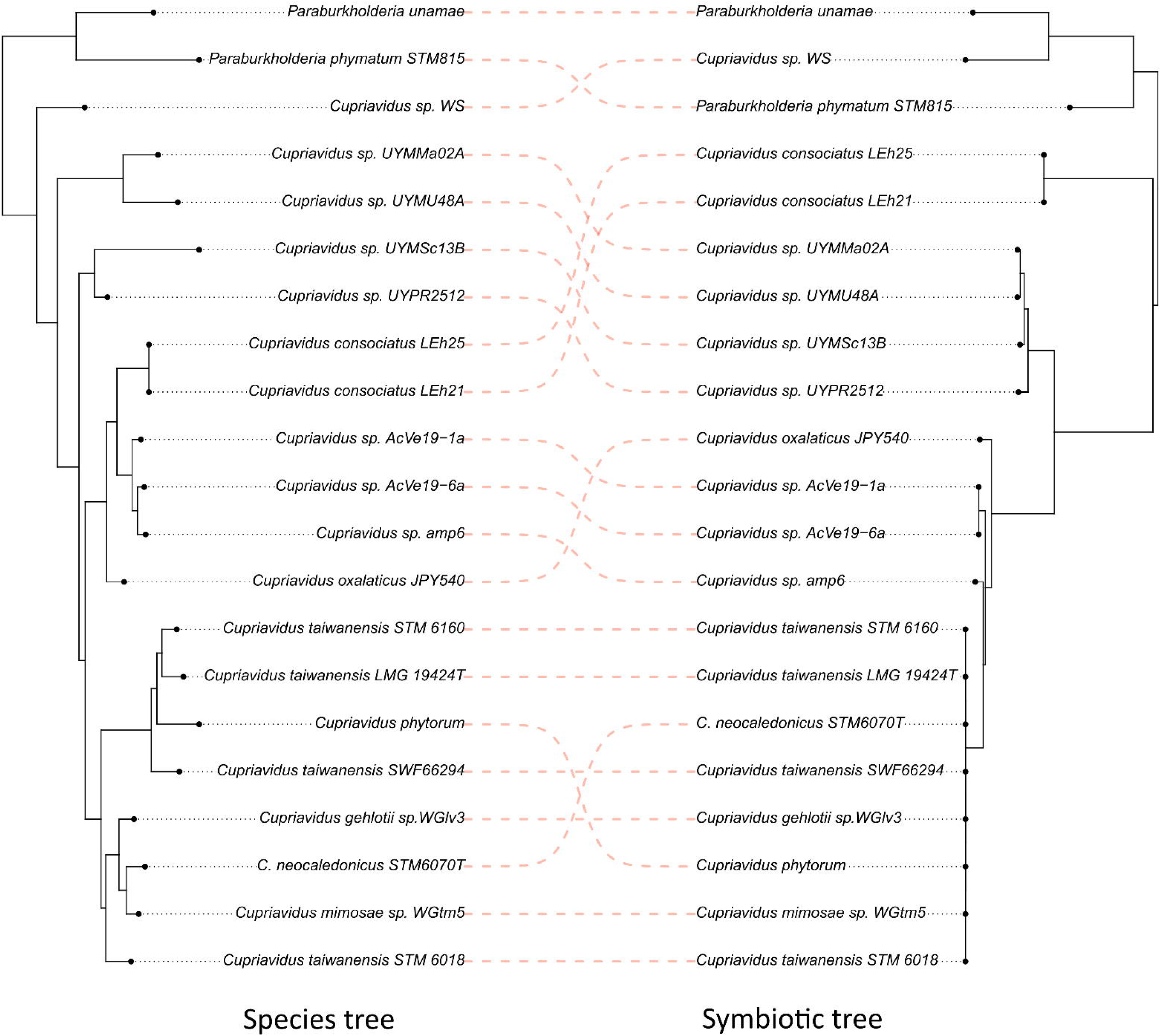
A) Phylogenetic relationships within the genus *Cupriavidus* and distribution of symbiotic genes. Maximum-likelihood phylogeny was reconstructed from core-genome alignments of representative strains of *Cupriavidus*, using Paraburkholderia as outgroup. **B)** The scheme on the right shows the presence/absence and percentage of identity of key symbiotic genes compared with UYPR2512, as reference. Tip labels highlighted in pink correspond to species and strains for which experimental evidence of nodulation has been reported. This figure illustrates the phylogenetic distribution of symbiotic traits within the genus.

In contrast, *C. ariasiae* UYMMa02A formed a separate lineage, closely related but distinct from *C. pinatubonensis* HN-2 and *C. yeoncheonensis* LMG 31506. This topology, together with low genomic similarity to *C. laharis*, supports the classification of *C. ariasiae* UYMMa02A as a novel *Cupriavidus* species. Previous phylogenetic analyses based on housekeeping genes showed that strain AMP6 represents an early-diverging lineage among nodule-forming *Cupriavidus* (Andam et al., 2007b; Andrus et al., 2012). However, our current phylogenomic analysis indicates that the branch containing strain UYMMa02A occupies a more basal position within the symbiotic *Cupriavidus* clade. This suggests that UYMMa02A represents the earliest diverging lineage of β-rhizobia within the genus.

Our results support the presence of at least five lineages containing symbiotic β-rhizobia *Cupriavidus*, in which rhizobial strains show a patchy phylogenetic distribution. The first lineage encompasses the well-known *C. taiwanensis* clade, including *C. taiwanensis* LMG19424^T^,*C. taiwanensis* STM 6160, *C. taiwanensis* SWF66294, and “*C. phytorum*” with the non-symbiotic species *C. nantongensis* X1^T^ and *C. nantongensis* HB4B5.

The second lineage is represented by *C. gehlotii* WGlv3, *C. mimosae* WGtm5, *Cupriavidus* sp. SS-3 with *C. neocaledonicus* STM6070^T^ and *C. neocaledonicus* STM6018, all of which carry symbiotic gene clusters. The third lineage contains the symbiotic strains *Cupriavidus sp.* UYPR2.512 and *Cupriavidus*. sp. UYMSc13B, which clusters with *C. necator* strains H850 and NBRC 102504, none of which have been reported as nodulating. The fourth lineage includes *Cupriavidus* sp. AMP6, “*C. consociatu*s LEh25”, and strains AcVe19-1a and AcVe19-6a, together with the non-symbiotic species *C. oxalaticus* NBRC 13593^T^. *C. oxalaticus* JPY540 is also present in this group and has been reported to nodulate *Mimosa pudica*, although its taxonomic assignment to *C. oxalaticu*s is uncertain (Rouws et al., 2024). Finally, the fifth lineage groups *C. ariasiae* UYMMa02A and *Cupriavidus* UYMU48A, whose closest sequenced relative is the non-symbiotic *C. pinatubonensis* HN-2.

The comprehensive analysis of all available *Cupriavidus* genomes also revealed additional lineages that may correspond to novel species and highlighted several cases of strain misidentification in public databases. Overall, our results demonstrate that symbiotic β-rhizobia within *Cupriavidus* are more diverse and phylogenetically widespread than previously recognized. However, it is noteworthy that the nodulation capacity appears to be restricted to a main lineage within the genus (spanning from *C. laharis* to *C. oxalaticus* clades), where it displays a patchy distribution. This pattern suggests that the symbiotic trait can be acquired by various lineages within this group and, in some species, may even behave as a polymorphism. This implies that species within this main lineage maintain a permissive genomic context for the acquisition and functional expression of the symbiotic plasmid, although the specific genomic elements enabling this compatibility are currently under study.

### Origin and organization of symbiotic genes

Given that *Cupriavidus* lineages within the beta rhizobia comprise both symbiotic and non-symbiotic strains and species, we were intrigued by the possible origins and evolutionary history of symbiotic genes in this genus. In alpha rhizobia, symbiotic capabilities are associated with the presence of *nif*, *nod*, and *fix* genes (Geddes et al., 2020; Lindström & Mousavi, 2020). To date, at least six *Cupriavidus* species are known to establish symbiotic associations with legumes of the genus *Mimosa*. Among them, *C. taiwanensis* has been reported as the main microsymbiont of widely distributed invasive *Mimosa* species, such as *M. diplotricha*, *M. pigra*, and *M. pudica* (W. M. Chen et al., 2001); *C. phytorum* isolated from the Zea mays and *M. diplotricha* rhizosphere in Mexico and Taiwan (Chávez-Ramírez et al., 2025), while *C. neocaledonicus* was obtained from *M. pudica* nodules growing in heavy-metal-rich mining soils (Klonowska et al., 2020). In Mexico, *Cupriavidus* strains, designated as “*C. consociatus* sp.” have been isolated from root nodules of *Leucaena* and *Arachis* species, respectively (Tapia-García et al., 2025). Although these strains carry key symbiotic genes, including *nifH*, *nifD*, and *nodC*, they are unable to fix nitrogen and form ineffective nodules, suggesting they are capable of initiating nodulation, but they do not establish functional nitrogen-fixing symbioses (Tapia-García et al., 2025). In *C. consociatus* (LEh25, LEh21), our re-annotation recovered a putative *nodB* homolog (∼90.7% identity) together with *nodA/nodC* (100/83.7% identity). However, the original description of *C. consociatus* reported the absence of *nodB* and ineffective (non-fixing) nodules on bean and *Leucaena* (white nodules). Since *nodB* (chitin oligosaccharide deacetylase) is essential for the synthesis of the Nod factor core structure, its presence explains the capacity of these strains to induce nodule formation. The observed phenotype (ineffective nodules) is therefore unlikely to be caused by the absence of *nodB.* An alternative and plausible explanation is that the natural host of these *C. consociatus* strains has not yet been identified. While Leucaena is considered a promiscuous legume capable of nodulating with diverse rhizobia, it is not a universal host. Thus, the symbiotic incompatibility observed (nodule formation without nitrogen fixation) may reflect host-specific constraints rather than a defective symbiotic machinery, suggesting that these strains might be effective symbionts of a different, yet unknown, legume partner. A conservative interpretation is that nodulation can be initiated (*nod* genes present), but nitrogen fixation fails, consistent with the phenotype described for these strains (Supplementary Note1). In the same study, two additional strains, AcVe19-1a and AcVe19-6a strains, were isolated and found to be positive for *nodC* amplification. Recently, these strains have been formally classified as part of the novel species *Cupriavidus phytohabitans* (Tapia-García et al., 2026). In line with this new taxonomic assignment, our genomic analysis reveals that both strains harbor a nearly complete symbiotic island, including nod, nif, and fix genes, with an organization comparable to other nodulating *Cupriavidus*. Consequently, we predict that C. phytohabitans strains AcVe19-1a and AcVe19-6a possess the genetic machinery required to induce functional nodules with appropriate plant hosts (Figure 2 A,B).

Recently, two novel species, ‘*Cupriavidus gehlotii*’ and ‘*Cupriavidus mimosae*’, have been proposed based on symbiotic isolates, with taxonomic descriptions currently available as preprints (Sharma et al., 2024). Notably, *Cupriavidus* sp. strain AMP6 shows exclusive association with *M. asperata* in North America (De Meyer et al., 2015). In South America, *Cupriavidus* sp. UYPR512 and diverse *Cupriavidus* spp. have been reported as effective symbionts of several native *Mimosa* species in Uruguay (Platero et al., 2016). The phylogeny generated also allowed us to determine the exact phylogenetic position of these nodule-forming *Cupriavidus* strains (Figure 2A).

The matrix displayed alongside the phylogenetic tree (Figure 2B) reveals the presence of symbiotic genes. Initial annotation suggested a ubiquitous presence of *nodI* and *nodJ* across the genus; however, genomic context analysis demonstrated that hits in non-nodulating strains correspond to ancestral paralogs lacking synteny with the nod operon (Aoki et al., 2013). Consequently, the authentic symbiotic *nodI* and *nodJ* orthologs are restricted exclusively to the symbiotic clades (Figure 2). Rather than being restricted to a single monophyletic lineage, symbiotic genes appear scattered throughout the tree. This patchy distribution suggests a complex evolutionary history likely shaped by a combination of horizontal gene transfer events and lineage-specific gene losses following an ancestral acquisition, rather than multiple independent origins.

Once the strains carrying symbiotic islands were selected, we concatenated their island genes and inferred a maximum-likelihood phylogeny (Figure 3A). The resulting tree recovered the five symbiotic lineages observed in the species tree, with most clades showing very high support (Boot ≥90). Nevertheless, some internal nodes displayed weak support (Boot <50), suggesting topological uncertainty and highlighting potential gene tree conflicts within the symbiotic region. We subsequently decided to compare not only the presence but also the organization of these symbiotic genes in some of the beta rhizobia lineages of *Cupriavidus* retrieved. We used the genetic organization of the *nif/fix/nod* cluster of UYMMA02A as reference for comparative analyses (Figure 3B). As mentioned before, the genes related to symbiosis in the chosen strains showed almost identical genetic composition and organization, despite the occurrence of minor differences mainly associated with insertions of transposase and other non-symbiotic-related genes. Consistent with this, the genomes contain the same 10 *nod* genes, arranged in a single cluster with the transcriptional regulator *nodD* located in opposite orientation to the other nine *nod* genes (*nodBCIJHASUQ*). This shared genetic organization, the unique copy of each gene, and the high degree of similarity between the lineages support the idea that these β-rhizobia synthesize the same *Nod* factors described for *C. taiwanensis*: pentameric chito-oligomers sulfated at the reducing end and N-acylated at the non-reducing end, frequently bearing an N-methyl and a carbamoyl group at the terminal residue (Amadou et al., 2008). This observation is coherent with the fact that all four strains are able to form effective nodules when inoculated on *Mimosa pudica* (Clerissi et al., 2018; Iriarte et al., 2016; Rouws et al., 2024; Vandamme et al., 2002). One of the most striking differences among the analyzed symbiotic islands lies in the organization of the nitrogenase *nif* operon. In the *C. taiwanensis* strain LMG19424^T^, we detected the insertion of four genes between *nifA* and *nifE*, arranged in the order pseudogene–transposase–transposase–pseudogene. The same genomic arrangement is also found in *C. taiwanensis* SWF66294 and *C.taiwanensis* STM 6160 (Figure S2). This structural disruption, already noted by (Amadou et al., 2008), has been proposed to influence the nitrogen fixation process. Interestingly, in *C. oxalaticus* JPY540, we also observe a separation between *nifA* and *nifE*, but in this case consisting of two transposases unrelated to those present in the *C. taiwanensis* strains. These observations suggest that the *nifA–nifE* intergenic region may represent a variable site where mobile elements recurrently insert, generating independent disruptions of the operon. In contrast, no interruption is observed in *Cupriavidus* sp. AMP6, *C. ariasiae* UYMMa02A, and *Cupriavidus* sp. UYPR2.512, as well as in other mimosoid-nodulating β-rhizobia, where the canonical *nifA–nifE* arrangement is conserved. Another relevant difference is observed within the nitrogenase operon organization. While most lineages conserve the canonical gene set, the symbiotic island of *C. ariasiae* UYMMa02A lacks a conventional *nifV* gene. Instead, sequence analysis revealed a gene fusion event between *nifV* and the downstream gene *nifW*, likely resulting from a mutation that disrupted the nifV stop codon (Figure S3). Interestingly, this chimeric fusion is conserved in other closely related strain (i.e., *Cupriavidus* sp. UYMU48A), a recurrence that excludes the possibility of sequencing errors and supports an ancestral fusion event in this lineage. This genomic feature aligns with the symbiotic ecology of the species; although *NifV* is essential for nitrogen fixation in free-living conditions and symbiosis with *Papilionoid* legumes, it has been experimentally demonstrated to be dispensable in *Mimosa* symbioses. In these interactions, the host plant is capable of supplying sufficient homocitrate to the bacteroids to sustain nitrogenase activity, thereby relaxing the selective pressure on the bacterial *nifV* gene (Bellés-Sancho et al., 2021; Hakoyama et al., 2009; Hashimoto et al., 2019; Lindström & Mousavi, 2020). Regarding the overall structure, the symbiotic island of *C. ariasiae* UYMMa02A spans 33.6 kb, with its length determined from a fully resolved single contig (Figure 3B). This confirms that, despite the specific remodeling of the *nif* operon (i.e., the *nifV*-*nifW* fusion), the island maintains the highly compact architecture characteristic of the *Cupriavidus* genus, comparable to the ∼35 kb island originally described for *C. taiwanensis* (Fig. 3B). This observations, together with the unique copy of each remaining gene and the high degree of similarity among lineages, supports the idea that this β-rhizobia are capable of producing *Nod* factors similar to those described for *C. taiwanensis* (Amadou et al., 2008), while displaying some degree of variability in the auxiliary genes of the *nif* cluster.

**Figure 3.**
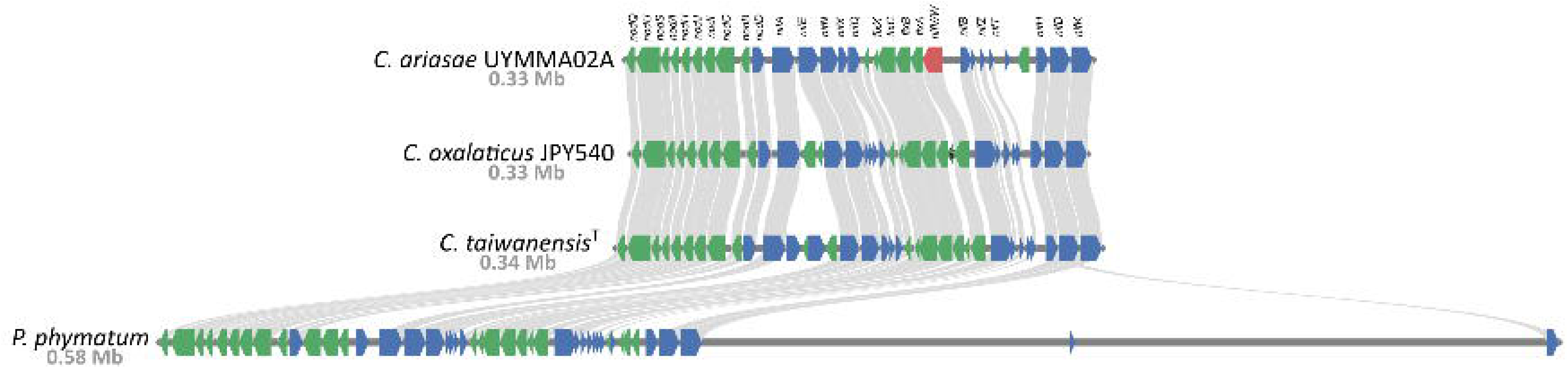
Evolutionary relationships and synteny of symbiotic islands in Cupriavidus. **(A)** Phylogenetic tree reconstructed from the cds sequences of symbiotic islands, showing their clustering as a monophyletic group within *Cupriavidus*. **(B)** Synteny analysis of symbiotic islands across representative Cupriavidus genomes, highlighting the conservation of gene order and the presence of locally collinear blocks. Synteny analysis of symbiotic islands across representative *Cupriavidus* genomes. Note that the *nifV*-*nifW* locus in *C. ariasiae* UYMMa02A is highlighted in red to indicate the chimera fusion. This chimeric open reading frame displays homology to both the independent *nifW* genes conserved in the other lineages, illustrating the structural divergence specific to the *C. ariasiae* strain.

Following, we can visualize that the symbiotic gene-based trees are not reflecting the same phylogenetic distribution observed when using core genomic information (Figure 4). In genome-wide based phylogenetic analyses, the *C. taiwanensis* clade appears closely related to the necator lineage, while *Cupriavidus* sp. UYMU48A and *C. ariasiae* UYMMa02A occupied basal positions. Conversely sym genes phylogeny cluster the *C. taiwanensis* species with the distant *Cupriavidus* sp. AMP6 strain, the two distantly related South American isolates together (AcVe19-6a, AcVe19-1a), and *C. oxalaticus* JPY540. Remarkably, the four Uruguayan isolates (*C. ariasiae* UYMMa02A, *Cupriavidus* sp. UYMU48A, *Cupriavidus* sp. UYMSc13B, and *Cupriavidus* sp. UYPR2.512) form a strongly supported monophyletic cluster in the symbiotic island-based phylogeny (Figure 3A), while they appear scattered across distinct lineages in the genome-wide phylogeny (Figure 2A). This incongruence strongly suggests that these strains, although genomically distant, harbor highly similar symbiotic islands. One possible explanation for the incongruent distribution of the symbiotic island and the core genome phylogenies is that the symbiotic trait, although ultimately derived from a common ancestor, has subsequently followed independent evolutionary trajectories in different *Cupriavidus* lineages. In addition, the high degree of synteny among *nod*, *nif*, and *fix* genes strongly suggests a shared origin for the symbiotic island. However, the phylogenetic reconstruction reveals contrasting evolutionary histories driven by geography and host ecology. First, we observe strong signals of regional endemism. The Uruguayan isolates form a robust monophyletic clade (Bootstrap = 100%) distinct from other lineages, suggesting a localized co-evolution in South America. Similarly, *C. consociatus* strains (LEh21, LEh25), isolated in Mexico, form a separate, highly supported cluster. This pattern is consistent with allopatric divergence, where native symbiotic populations in geographically distant regions (South vs. North/Central America) have evolved independently over time. In contrast, the clade containing *C. taiwanensis*, *C. neocaledonicus*, and *C. mimosae* displays a different pattern. While the broader grouping is supported, the internal nodes show extremely low resolution (Bootstrap < 50%), indicating that these symbiotic islands share near-identical sequences despite being isolated from disparate regions (e.g., Taiwan, New Caledonia, Central America). This lack of genetic resolution is consistent with a recent, rapid global dissemination rather than ancient historical proximity. The presence of *C. neocaledonicus*, originally described in the Pacific, within this largely American/Asian cluster is likely explained by the invasive nature of their hosts. Since *Mimosa* species (e.g., *M. pudica*, *M. diplotricha*) are aggressive invaders introduced globally from the Americas, they likely acted as vectors, transporting their symbiotic islands across continents. Thus, the grouping of these geographically diverse strains likely reflects anthropogenic sympatry driven by biological invasion.

**Figure 4.**
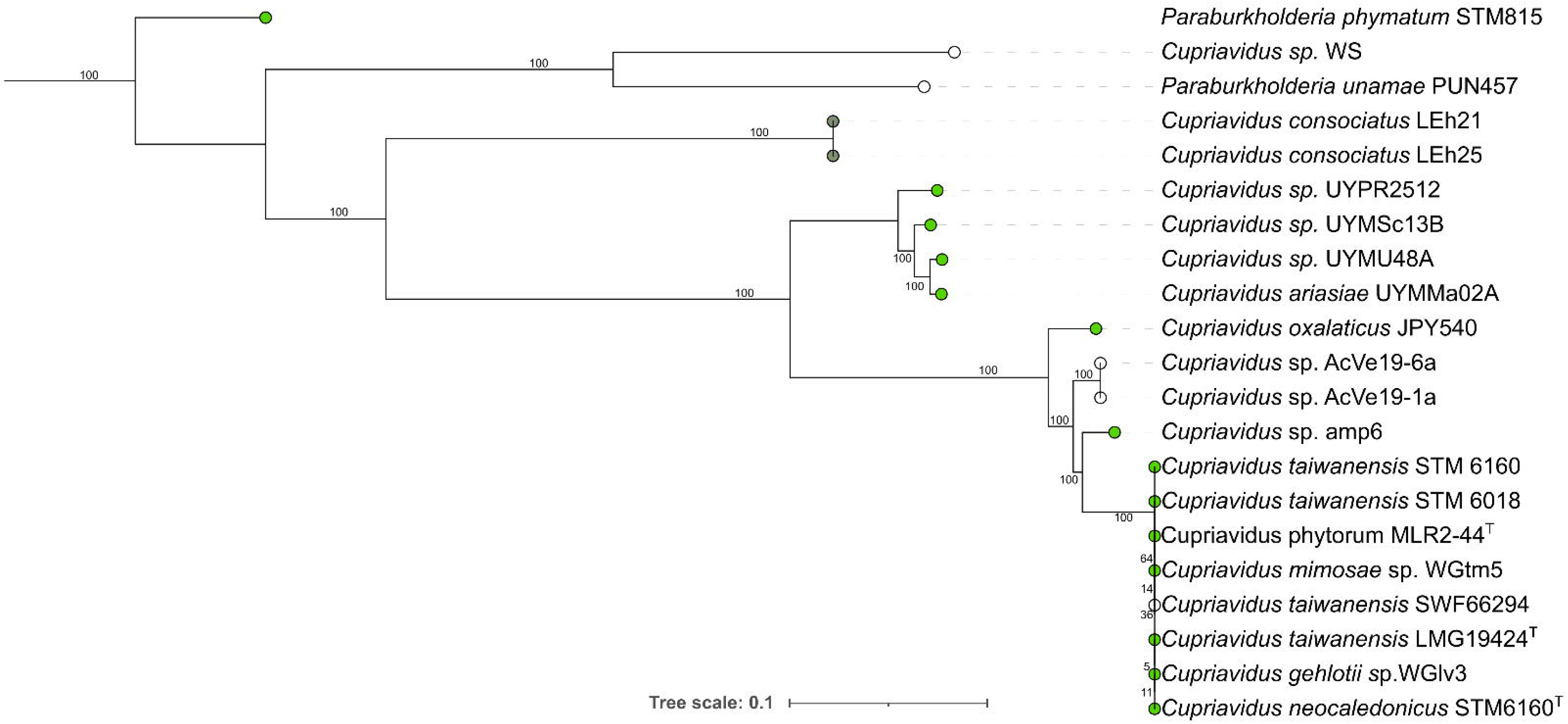
Tanglegram comparing the phylogenomic species tree and the symbiotic island phylogeny of *Cupriavidus* nodulating strains. (Left) The species tree was reconstructed using PhyloPhlAn 3.0 based on a concatenated alignment of 400 universal conserved marker genes. (Right) The symbiotic phylogeny was inferred from a concatenated alignment of key symbiotic genes (nodA, nodC, nifH, fixA) located within the symbiosis island. Both trees were inferred using Maximum Likelihood in IQ-TREE with 1000 bootstrap replicates. Connecting lines link the same strain between the two phylogenies. Crossing lines indicate incongruence between the core genome and the symbiotic island.

Thus, both ancestral conservation and local horizontal gene transfer appear to have shaped the current distribution of symbiotic traits within the genus. Supporting this hypothesis, previous work by our group showed that partial *nifH* and *nodA* sequences from Uruguayan *Mimosa*-nodulating isolates clustered together with Brazilian *C. necator* symbiotic isolates, forming a separate branch from the *C. taiwanensis* clade (Platero et al., 2016). Also, the analysis of five concatenated sym plasmid loci from 80 Caribbean *Cupriavidus* symbiotic isolates showed that they formed a monophyletic clade (haplotypes A–E), clearly separated from *Cupriavidus* sp. UYPR2.512, the only South American strain included in the study (Parker, 2015).

Overall, the results presented in this work show that all known β-rhizobial *Cupriavidus* strains share almost the same set of symbiotic genes, organized in a few equally arranged operons with a high level of synteny. This highly conserved ordering of sym genes across distinct *Cupriavidus* lineages suggests that purifying selection may act to preserve their genomic arrangement, reflecting functional constraints in the regulation and expression of the symbiotic machinery. These observations support the hypothesis of a single ancestral acquisition of symbiotic abilities within the genus, most likely from a β-rhizobial Paraburkholderia, followed by the subsequent loss of these abilities in some species within each lineage (Amadou et al., 2008; W.-M. Chen et al., 2005). Nevertheless, we cannot rule out that the phylogenetic distribution of sym genes observed in *Cupriavidus* may also be explained by horizontal gene transfer events among species from related lineages sharing common environments.

### Comparative genomics of symbiotic plasmids in *Cupriavidus*

One of the main discoveries in the study of nodulating beta Proteobacteria was that genes directly responsible for the symbiotic and nitrogen fixation phenotype are located in plasmids (W.-M. Chen et al., 2003). This was also stated in many species of alpha-rhizobia (Yang et al., 2020). Recent works have addressed the question of whether the symbiotic plasmid in beta-rhizobia is the same one in all species (Black et al., 2012; Parker, 2015; Zheng et al., 2017).

Based on phylogenetic and sequence analysis (Parker, 2015) proposed that sym plasmid inheritance in *Cupriavidus* is mainly vertical. Nucleotide diversity estimation based on *nodA*, *nodC*, *nifH*, *fixA*, and *phaZ* showed that these genes display lower nucleotide diversity than chromosomal housekeeping loci, and that they cluster as a monophyletic group within *Cupriavidus*. Interestingly, since all but *phaZ* are located within the symbiotic island, this pattern likely reflects the evolutionary trajectory of the island itself rather than that of the plasmid or the core genome. This observation highlights the symbiotic island as a cohesive evolutionary unit, whose dynamics may obscure signals from the broader genomic background.

We investigated the conservation of symbiotic plasmids within the Burkholderiaceae family. Nucleotide sequence identity among complete plasmids from Beta-rhizobium genomes was evaluated using the BLAST+ suite. This analysis, using the symbiotic plasmid of *C. taiwanensis* LMG19424^T^ as a query, revealed two distinct patterns of conservation across the Burkholderiaceae. Only six strains, all belonging to *C. taiwanensis*, displayed nearly complete coverage of the plasmid, consistent with the report of Clerissi et al. (2018) (Clerissi et al., 2018) that this plasmid (pRalta) was acquired once in the ancestor of *C. taiwanensis* and has since remained highly conserved. In contrast, all other *Cupriavidus* species exhibited very limited similarity (Table S3). We were curious as to whether this restricted homology was simply due to the presence of symbiotic genes, and therefore intersected the BLAST alignments with the coordinates of the symbiotic island (see Methods). To quantify the conservation of the symbiotic plasmid across Burkholderiaceae, we computed the coverage of the reference plasmid (pRalta, *C. taiwanensis* LMG19424) by each subject sequence. Coverage metrics are summarized in Table 1 and Figure 5. For each subject, we report the total number of nucleotides aligned to the plasmid, the nucleotides mapping within the symbiotic island, and those mapping outside of it. Percentages are expressed relative to the plasmid (for total, island, and backbone coverage) and relative to the island alone (percentage of the island covered).

**Figure 5.**
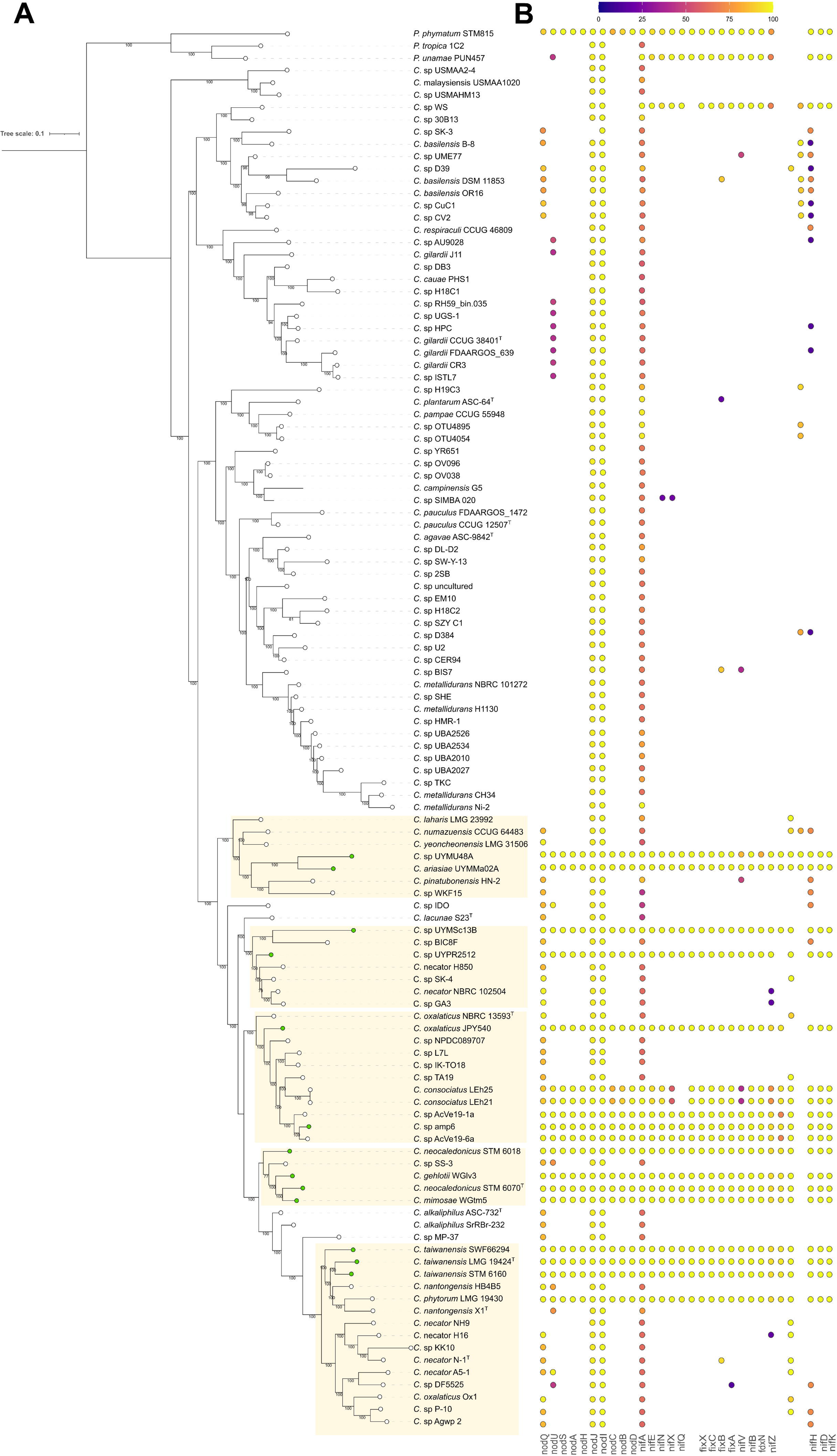
Conservation patterns of the symbiotic plasmid pRalta across Burkholderiaceae. BlastN hits from selected *Cupriavidus* genomes were mapped against the reference symbiotic plasmid pRalta from *C. taiwanensis* LMG19424^T^ (GenBank: NC_010529.1). The x-axis represents the coordinate position (bp) on the reference plasmid. The shaded vertical band marks the location of the symbiotic island (approx. 302–337 kb). Horizontal bars indicate regions of homology: orange segments correspond to matches within the symbiotic island, while blue segments represent matches to the plasmid backbone.

**Table 1.** Conservation of the reference symbiotic plasmid pRalta (*C. taiwanensis* LMG19424^T^) across Burkholderiaceae plasmids. BLASTn alignments were used to compute the total number of nucleotides matching the reference plasmid (nt_total), as well as the fraction of alignments located inside the symbiotic island (nt_in_island) or in the plasmid backbone (nt_out_island). Percentages are expressed relative to the plasmid as a whole (pct_total, pct_isla_plasmid, pct_out_plasmid) and to the island alone (pct_isla_relative).

While many partial matches indeed mapped to *nod* and *nif* loci, resulting in high relative coverage of the symbiotic island, several subject plasmids also displayed substantial similarity outside this region. This indicates that, although the symbiotic island explains a large fraction of the similarity, certain conserved modules from the plasmid backbone also contribute to the observed matches. Despite these shared modules, the overall plasmid backbones are highly divergent, revealing a mosaic structure. This suggests that the symbiotic island is an independent mobile element capable of horizontal gene transfer among plasmids of different origins. Such mobility is consistent with the high number of insertion sequences and transposases flanking the island.

Searches for homologous genes to pRALTA were also done in all *Cupriavidus* assemblies, including draft genomes. Again, we identified homologous symbiotic genes among the rhizobial genomes, supporting that the island is highly conserved in *Cupriavidus*, while the rest of the plasmidic genes are only marginally conserved.

Taken together, our results support the high conservation of symbiotic genes across beta-rhizobia, a pattern consistent with purifying selection acting on their sequence evolution. However, the presence of sym genes is not completely fixed in the species, suggesting that their distribution may behave as a polymorphism. In addition, the observed conservation patterns indicate that the symbiotic plasmid may be evolving in a mosaic fashion. This could involve integration of genes from other plasmids or mobilization of the symbiotic island into new plasmid backbones. In other words, beyond vertical inheritance, hybridization and horizontal gene transfer among plasmids from closely related strains may also explain the current structure of beta-rhizobial sym plasmids (Zheng et al., 2017). Similar processes have been proposed for alpha-rhizobial sym plasmids as well (Reeve et al., 2010).

## CONCLUSION

Our comparative genomic analyses reveal that symbiotic *Cupriavidus* species are more diverse than previously recognized, encompassing at least five independent lineages intermixed with non-symbiotic species. Within these lineages, the arrangement and sequence conservation of the *nod*, *nif*, and *fix* operons remain remarkably stable, supporting the hypothesis of a single ancestral acquisition event of symbiotic capacity, followed by secondary losses and horizontal transfers. However, the comparative analysis of plasmid sequences shows that conservation is largely restricted to the symbiotic island, whereas the plasmid backbone displays a mosaic structure, consistent with recurrent recombination and integration of mobile elements. This dual pattern, strong purifying selection acting on the symbiotic operons and high plasticity of their plasmid carriers, explains both the widespread distribution of symbiotic genes across distinct *Cupriavidus* clades and their absence in some closely related strains. Altogether, our results underscore the complex evolutionary history of β-rhizobia within *Cupriavidus*, where vertical inheritance, polymorphic distribution of symbiotic genes, and horizontal gene transfer collectively shaped the current diversity of symbiotic plasmids. These findings expand the evolutionary framework of legume–rhizobia interactions and suggest that novel symbiotic *Cupriavidus* species will likely continue to be discovered.

## Data availability statement

The genomic sequences analyzed in this study are publicly available in the NCBI database. The specific accession numbers for the genomes utilized for phylogenetic reconstructions and comparative analyses are listed in Supplementary Table S1. Additionally, the custom scripts and computational pipeline used for the plasmid conservation metrics and comparative genomics are openly available in the GitHub repository: https://github.com/Melisa-Magallanes/Plasmid-analysis.

## Ethics Statement

This article does not contain any studies involving human participants or animal subjects performed by any of the authors. All experimental procedures involving plant nodulation assays were carried out in accordance with standard institutional guidelines and local regulations.

## Acknowledgments

The authors’ work was supported by PEDECIBA and ANII-Uruguay FCE_1_2014_1_104338 to RP, ANII-Uruguay FCE_3_2013_1_100727 to AI. JS, AI, and RP are members of the PEDECIBA and Sistema Nacional de Investigadores, Uruguay.

## Credit authorship contribution statement

Melisa Magallanes Alba: Investigation; Formal analysis; Visualization; Writing – original draft. Cecilia Rodríguez-Esperón: Investigation; Methodology.

José Sotelo-Silveira: Software; Formal analysis; Resources.

Elena Fabiano: Conceptualization; Funding acquisition; Writing – review & editing.

Andrés Iriarte: Conceptualization; Software; Formal analysis; Supervision; Writing – review & editing.

Raúl Platero: Conceptualization; Project administration; Funding acquisition; Supervision; Writing – original draft; Writing – review & editing.

---

**Supplementary Figure 1.**
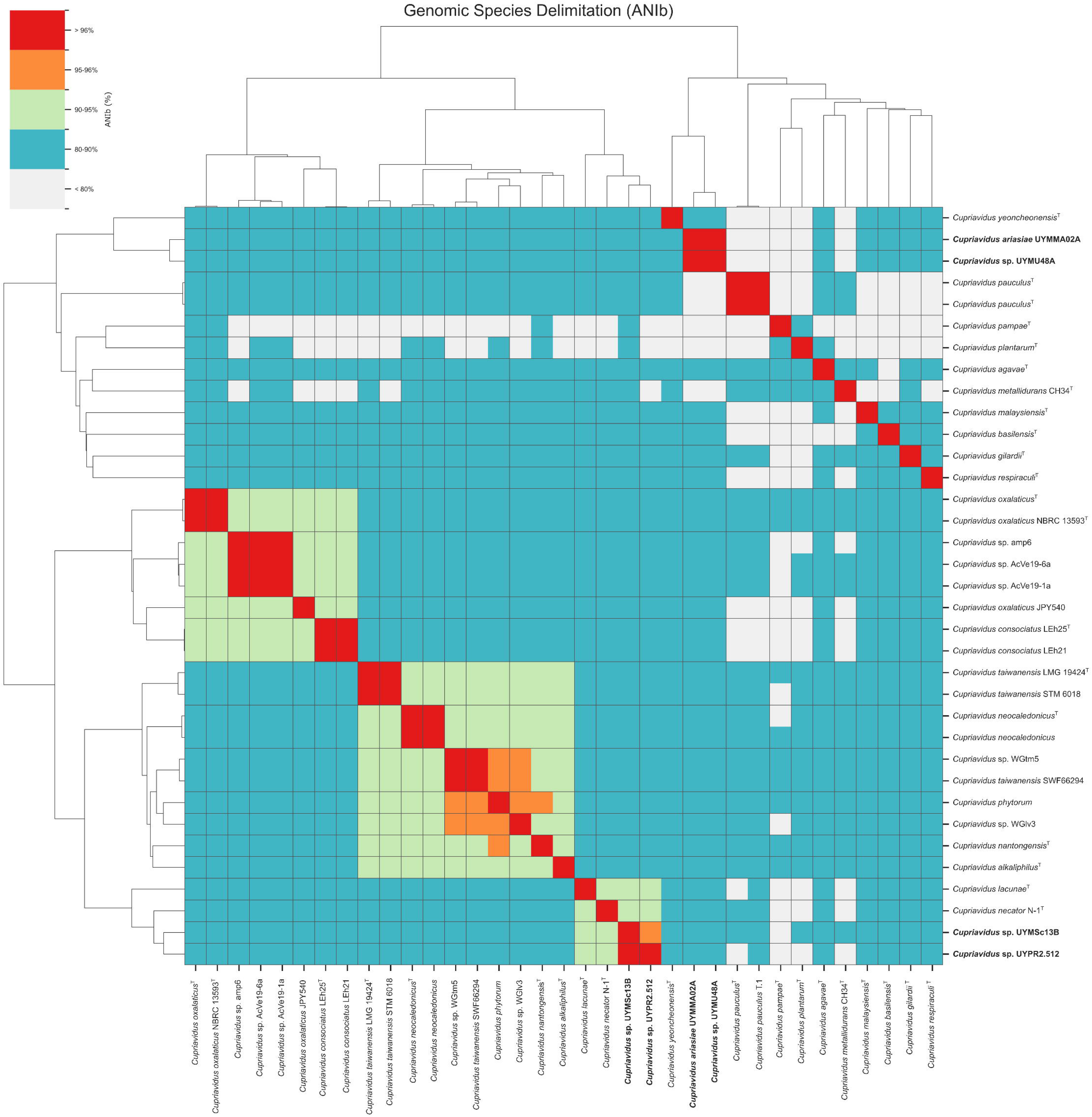
Comprehensive genomic relatedness among all studied *Cupriavidus* strains. Pairwise genomic comparison matrices for the complete dataset (n = 114 genomes), visualized as hierarchically clustered heatmaps. The analysis employs three complementary metrics: **(A)** Average Nucleotide Identity (ANIb) calculated via BLAST+; **(B)** FastANI, an alignment-free ANI approximation; and **(C)** Estimated digital DNA–DNA hybridization (dDDH), derived from ANIb values using a Generalized Linear Model (GLM). Strains are ordered according to the associated dendrograms (Euclidean distance, average linkage). Color scales are discretized to highlight species boundaries: red indicates species identity (ANI >96%, dDDH >70%); orange denotes the speciation threshold zone (ANI 95–96%, dDDH 50–70%); and blue/grey represents distinct species (ANI <90%, dDDH <50%). This global analysis identifies *C. necator* H850 as the closest genomic match to *Cupriavidus* sp. UYPR2.512 (ANIb ∼94.8%, dDDH 60.2%), yet confirms that the strain remains below the species cutoff. Similarly, *C. ariasiae* UYMMA02A exhibits low similarity to all validly described species (ANIb <89%, dDDH <38%), supporting its classification as a novel species.

**Supplementary Figure 2.**
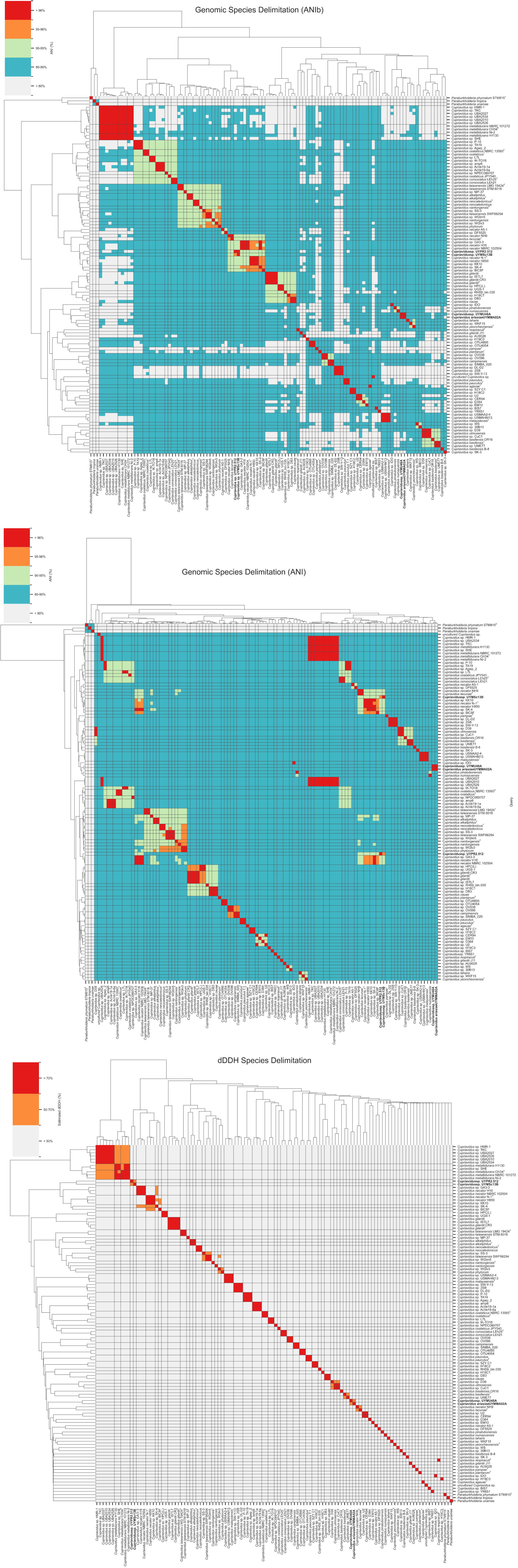
**Insertion of transposases between nifA and nifE in three *Cupriavidus taiwanensis* strains**. Comparative genomic analysis revealed the insertion of four genes between nifA and nifE in C. taiwanensis LMG19424^T^. These genes are arranged in the order pseudogene–transposase–transposase–pseudogene. The same genomic arrangement is also present in *C. taiwanensis* SWF66294 and *C. taiwanensis* STM6160, indicating a species-level conserved structural feature across these strains that may reflect shared mobilization events in the symbiotic plasmid.

**Supplementary Figure 3.** Domain architecture of the chimeric nifV-nifW locus in *C. ariasiae* UYMMa02A. InterProScan analysis identifies Homocitrate synthase domains at the N-terminus (Pfam PF00682: HMGL-like, residues 1–212) and NifW domains at the C-terminus (Pfam PF03206: Nitrogen fixation protein NifW, residues 368–463).

**Supplementary Table 1.** List of genomes used in the present work.

**Supplementary Table 2.** Comprehensive pairwise genomic comparison matrices for the genus Cupriavidus. The file contains three spreadsheets providing the complete all-vs-all similarity values for the 114 genomes analyzed in this study: ANIb (Average Nucleotide Identity based on BLAST+) calculated using PyANI, FastANI (Whole-genome ANI approximations calculated using FastANI) and Estimated dDDH (Digital DNA–DNA hybridization (dDDH) values estimated from ANIb data using a Generalized Linear Model (GLM). Values are expressed as percentages.

**Supplementary Table 3.** Conservation of the *C. taiwanensis* symbiotic plasmid across Burkholderiaceae genomes. . Coverage of the reference plasmid pRalta (strain LMG19424^T^) was assessed by BLASTn against complete plasmids from Beta-rhizobium genomes. Two main patterns were observed: (i) nearly complete coverage in six *C. taiwanensis* strains, consistent with a single ancestral acquisition followed by strong conservation, and (ii) very limited similarity in other Cupriavidus species, where matches were largely restricted to the symbiotic island.

